# Reply: Neutral tumor evolution?

**DOI:** 10.1101/274142

**Authors:** Timon Heide, Luis Zapata, Marc J. Williams, Benjamin Werner, Chris P. Barnes, Trevor A. Graham, Andrea Sottoriva

**Author notes:** These authors contributed equally to this work (Timon Heide, Luis Zapata, Marc J. Williams, Benjamin Werner).

## Abstract

Mutation, selection and neutral drift shape the cancer evolutionary process^1^. The role of selection has received particular interest, but inferring the presence and strength of selection during tumour growth remains challenging. Recently, we analysed the frequency distribution of subclonal mutations in many cancers and found that in approximately 30% of cases the observed distribution was entirely consistent with a simple model of neutral evolution^2^. Thus, we concluded that neutral evolution, perhaps surprisingly, provides an adequate explanation of the intra-tumour heterogeneity present in a significant proportion of cancers.

Tarabichi and colleagues [bioRxiv: 2017/06/30/158006] question the robustness of the method we presented in Williams *et al*. 2016^2^ to identify neutral cancer evolution from variant allele frequency (VAF) distributions. Their critique has four main points that we address in this document using a simulation approach and a reanalysis of public datasets.

## 1. Impact of clonal copy number alterations

In Williams *et al*. 2016^2^, we assessed the cumulative VAF distribution M(f) over the frequency range of f=[0.12,0.24] in order to restrict our analysis to subclonal variants. We classified a tumour as non-neutral if M(f) was not proportional to 1/f in this range.

Tarabachi and colleagues note that tumours with a tetraploid genome will have a ‘peak’ of clonal mutations at f~0.25 (clonal mutations in a single allele), thus causing an ‘artificial deviation from the linear fit’ and hence incorrect rejection of neutrality.

First, we note that in cases where cancer growth is *initiated* by a genome doubling, there will be no peak at f=0.25, and a 1/f neutral tail will still be produced with a coefficient twice the effective mutation rate. Indeed, the variable timing of genome doubling relative to the initiation of tumour growth, and its impact on the VAF distribution, is one of the key confounding factors in calling a cancer genome as tetraploid^4^.

Second, the important confounding issue of ploidy concerned us in the original manuscript. Assuming a tumour with ploidy π and a clonal peak that is binomially distributed, the expected standard deviation of the peak is 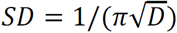, where D is the (mean) read depth. For the 1/f fit we aimed at excluding mutation frequencies above 1/π-2*SD. We chose our integration range based on a triploid tumour with read depth of 100X, giving an upper threshold of 0.26. Although this is suitable for the majority of cases, we agree that a tetraploid or higher ploidy tumour could have a peak within our integration range, and therefore potentially more tumours could be consistent with neutral evolution than those we reported.

In general, we do acknowledge that the 1/f integration method is sensitive to the choice of integration range, and that while that represents the correct analytical derivation, it has important limitations when applied to real data. To tackle these limitations, we recently developed more accurate metrics, as well as a Bayesian model selection framework that directly compares the neutral model against models with selection using *the entire VAF distribution*, thus avoiding the shortcomings of the 1/f analytical form^3^. We also contributed to the development of additional classification methods for neutrality that exploit multi-region sequencing^4^.

Since all bioinformatics methods have limitations, in both our original manuscript and follow-up studies we focused on both improving the methods as well as validating our findings in higher depth data (e.g. WGS gastric cancers^2^) and multi-region sequencing datasets^4^. We care to stress however, that the majority of cancers we analysed were *not neutral*, and showed signs of subclonal selection.

## 2. Interpretation of the 1/f statistical test

Tarabachi and colleagues correctly note that failing to reject the null is not necessarily evidence for the null. This is absolutely true, but evolutionary analysis of cancer genomic data requires a hypothesis-driven approach based on a sensible null. Analysing data without knowing what to expect in the simplest scenarios may lead to wrong conclusions, as we have recently highlighted^4^. The fundamental message in our original manuscript is that neutrality, the null model of molecular evolution^5^, was often a sufficient explanation of the available data, an entirely valid approach from a frequentist perspective^6^.

In their letter, the authors state that in order to infer neutrality one would have to reject all non-neutral models. We believe this is a fundamentally flawed position for two reasons. First, the test we applied quantifies the deviation from the null distribution in terms of a change in model parameter (s = 0 vs s > 0). Arguments for setting up the test in any other way seem arbitrary since no model, neutral or otherwise, could be considered a ‘true’ model. One case where tests arise is when the null is expected to be approximately true or there is strong interest in establishing that it is likely to be false^6^, which is exactly the case in point. Second, because there are an infinite number of models of selection, some selection models, such as those with very weak selection, produce tiny deviations from neutrality that are not measurable. Others are biologically unrealistic (e.g. each single mutation in the genome is a driver), or do not apply to cancer (e.g. constant population size). The infinite number of selection models is why, in molecular evolution, neutrality is the null^7^. In our view, selection remains arguably the most important force in cancer, leading to phenotypic adaptation, but maintaining a sensible null model is required to avoid over-interpreting the data.

Tarabichi and colleagues state that the deterministic solution we report in our manuscript (Eq.7) relies on the strong assumption of synchronous cell divisions. This statement is false, there is no such assumption in that analytical model, which in fact is the convergent solution of a continuous-time stochastic branching process^8^. Relatedly, while our simulations in the original manuscript did assume synchronous cell divisions, in our most recent pre-print work^3^ we do not make such an assumption, and show that asynchronous cell divisions do not alter the conclusions. A comprehensive analysis of the underlying stochastic Luria-Delbrück model shows that the scaling behaviour remains unchanged even in the explicit presence of stochastic cell death^9^. Therefore, for data of sufficient depth, ‘stochastic, effectively neutral evolution’ does imply and is implied by a 1/f cumulative distribution. In Tarabichi’s letter Figure 1b, the claim that a stochastic neutral model does not imply 1/f is generally false, as we showed in our original manuscript and as demonstrated by others before us^8–10^.

**Figure 1.**
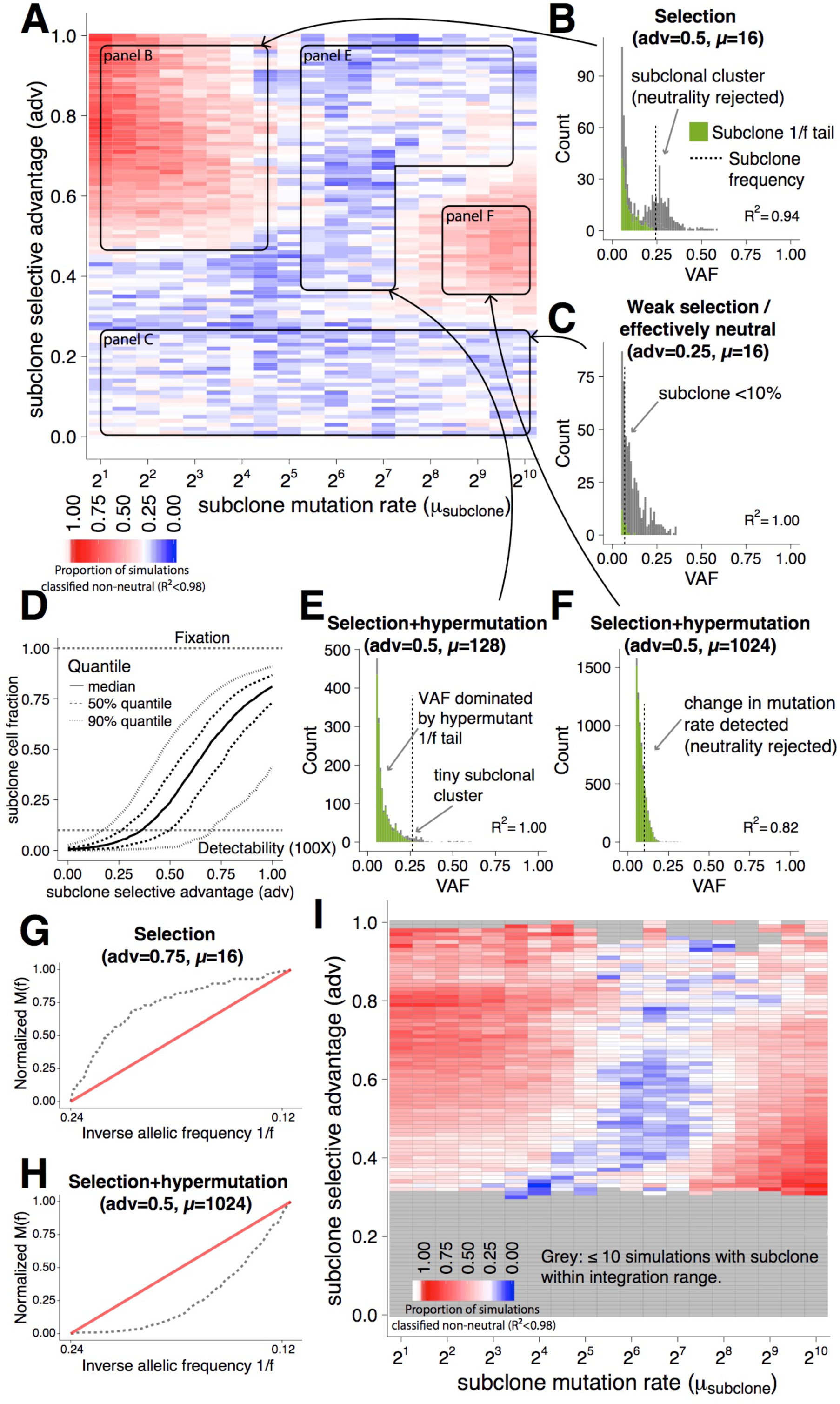
Insights from stochastic simulations of cancer growth. **(A)** Heatmap recapitulating Tarabichi’s Figure 1b showing proportion of simulations where neutrality is rejected as a function of subclone fitness advantage and mutation rate with parameter sets from Tarabichi’s letter. **(B)** Example VAF distribution with a detectable subclonal cluster (dashed line indicates subclone frequency) in which the 1/f test rejects neutrality in favour of selection (R^2^ value reported). **(C)** Example VAF distribution with a weakly-selected subclone that remains below the limit of detection in the data (100X depth). **(D)** Subclone cell fraction in the funal tumour as a function of fitness advantage, for adv_subclone_<0.5 the subclone rarely reaches the detectable size of ~10% cell fraction. **(E)** Example VAF distribution for a subclone with selective advantage and, at the same time, high mutation rate (and corresponding cumulative 1/f plot, panel **G**). **(F)** Example VAF distribution for a selected and extreme mutator subclone (and corresponding cumulative 1/f plot, panel **H**). **(I)** Heatmap showing proportion of simulations where neutrality is rejected for the subset of cases where the selected subclone had a final frequency within the integration range of [0.12,0.24] used in the 1/f test (signature of selection was within the 1/f test).

Tarabichi and co-authors state that assuming constant cell death and constant mutation and division rates is unrealistic. However, the authors then present a classical stochastic branching process simulation approach where these values *are* constant, and change only when selection comes to play (phenotypic change in either the selection coefficient of the mutation rate). That is *precisely* what we presented in the computational simulations in our original manuscript (see “In silico validation of the neutral model”, page 242 of Williams *et al*. 2016^2^). Hence, we believe this is not a valid criticism of our work.

In their letter, the authors state that simulating stochastic processes is more realistic than using the corresponding deterministic solution. We agree, and this is precisely the reason why in our original manuscript we used stochastic simulations that modelled random cell death and to show the convergence to the analytical form (Fig S9-S11). Tarabichi *et al*. refer to Bozic *et al*. 2016^11^ as an example of a better study. Bozic *et al*. 2016 is a sound and interesting study that appeared shortly after our Williams *et al*. 2016 manuscript and corroborates our results on neutrality, also citing the bioRxiv version of the Williams *et al*. paper (citation [34] in the Bozic *et al*. paper). Furthermore, the solution in the Bozic study is identical to the one we presented (Williams *et al*. Eq.7 vs Bozic *et al*. Eq.7), again confirming that the stochastic and deterministic solutions converge. We think this comment misrepresents our work and distracts from the core points of the discussion.

Tarabichi *et al*. state that the biological noise modelled in branching processes can lead to a deviation from the deterministic (convergent) solution. This is of course true, and indeed was the motivation for our original *in silico* validation (where we also simulated sequencing noise). We also note that a large component of the ‘biological noise’ the authors refer to is *genetic drift*, which can lead to the formation of subclonal clusters without selection (thus rejection of 1/f). However, genetic drift is relevant only in small populations, during the first few division of the growing tumour, as drift in large populations is very weak. We note that in a growing tumour, a small population is also a *young* population, and so not many mutations will have been accumulated in few divisions elapsed in each lineage, thus drift clusters are expected to contain only a very small number of mutations, likely not enough to cause a substantial deviation from 1/f. Even in the cases where drifting clones do lead to rejection of the 1/f fit, this leads to *underestimation* of neutrality using the frequentist test.

## 3. Insights from simulated tumours

Tarabichi and colleagues use a stochastic branching process, similar to the one we described in our original publication^2^ (online methods, Fig. S9-S11) and in our follow-up 2016 pre-print^3^, to generate synthetic genomic data and test the 1/f linear fit. Specifically, in their Figure 1, Tarabichi *et al*. present a synthetic analysis of the 1/f test using the analytical deterministic solution (Figure 1a) and stochastic simulations (Figure 1b). In both analyses, a new subclone is introduced at a certain fixed time point (e.g. at the 8^th^ generation in Figure 1b). The authors claim that the 1/f test does not discriminate between neutral and non-neutral tumours in their analysis.

First, we noted that the stochastic simulation result does not appear to converge to the deterministic one (i.e. Tarabichi’s Figure 1a is very different from 1b), which is at odds with previous literature^2,8–11^. Second, we were surprised by the choice of simulated parameters in their synthetic test. In the authors’ analysis, the driver event in the newly selected subclone does not just induce a selective advantage, modelled as an increase net-growth rate (adv_subclone_), but *at the same time*, also a change in mutation rate. Curiously, the new subclone can have lower mutation rate (up to 8 times lower than the background mutation rate of 16 mutations per division) or higher mutation rate (up to 100 times higher). A mutation rate of 1024 mutations per cell division (see Tarabichi’s Figure 1a, x-axis, *μ_subclone_* = 2^10^) is found only in a very small set of colorectal or uterine cancers characterised by mutations in the proof-reading domain of POLE or POLD. To our knowledge, a POLE subclone arising within a POLE wild-type background has never been documented. Thus, we urge caution when considering the implications of the parameters at the extremities of the range considered by Tarabichi *et al*.

In an attempt to understand the points of Tarabichi *et al*., we have reproduced Tarabichi’s Figure 1b using our stochastic branching process simulation with precisely the same parameters (Figure 1A in this document, see Methods for details). Our stochastic analysis did recapitulate the Tarabichi’s deterministic analysis in *their* Figure 1a, as one should expect^2,8–11^.

We found that when selection generated a detectable subclonal cluster with f_subclone_≥10%, this was correctly identified by the 1/f test in the majority of cases and neutrality was rejected (top left quadrant of Figure 1A, example in Figure 1B, 1/f tail of new subclone in green). For the majority of cases where the 1/f test failed, this was due to the new subclone being very small because of weak selection (adv_subclone_<0.5, bottom half of Figure 1A, example in Figure 1C). Figure 1D illustrates, using the same simulations as in Figure 1A, the relationship between selective advantage and the subclone cell fraction in the final tumour, highlighting the issue of weak selection. We have specifically quantified this effect in our 2016 pre-print^3^, identifying a ‘wedge of selection’ that describes this detectability problem in cancer genomic data at current resolution. In general, the fact that selection is inefficient in expanding populations is well known in population genetics^12^ and we have demonstrated this effect in colorectal cancer^5^. If subclonal selection does not significantly change the clonal composition of the tumour, the signature of neutral growth (a ‘1/f tail’) dominates the detectable VAF spectrum (Figure 1C). In these cases the tumour is driven by ‘effectively neutral’ dynamics, as we recently demonstrated using multi-region sequencing data from multiple cancer types^4^.

Notably, on the right-hand side of Figure 1A, where the subclone had both a selective advantage and was also hypermutant (*μ_subclone_*>=64), we found that interesting evolutionary dynamics were taking place. The analysis showed that a hypermutant subclone generates a massive 1/f tail containing thousands of the subclone’s own private mutations that dominate the entire VAF distribution, obscuring the underlying subclonal structure (which was generated with the old mutation rate, and hence with many less mutations). In the VAF distribution of these tumours it is very difficult to identify the subclonal cluster because it is tiny (it has only a few hundred mutations) with respect to the subclone’s enormous 1/f tail, generated with up to 1000 mutations per cell division (top right quadrant of Figure 1A, example in Figure 1E). It is not surprising that our test, or any other test, would struggle to detect any subclonal cluster or deviation from 1/f in these VAF distributions. In this potentially unrealistic scenario, we suggest however that Tarabichi and colleagues have clearly demonstrated how 1/f tails can be the dominant signal in cancer genomic data, to the extent where they can dominate the entire subclonal structure. Curiously, for moderate values of selection (adv_subclone_~0.5), a change in mutation rate from normal to hypermutant could be detected, leading to rejection of neutrality (mid-right area in Figure 1A and example in Figure 1F). An analogous deviation from 1/f caused by an increase in mutation rate was discussed in Williams *et al*. 2016, Figure S11H. Interestingly, rejection of the null was not caused by a bulging of the cumulative distribution upwards as in the case of selection (Figure 1G), where there are more mutations at high frequency than expected under neutrality, but instead pushed the distribution downwards (Figure 1H). For weak selection and a hypermutator subclone, the new subclone did not reach a detectable size and therefore neutrality could not be rejected (as in Figure 1C). These dynamics are interesting, although a subclone with 10–100 times higher mutation rate with respect to the background clone may be a rare event.

In some cases, the subclone may grow to a frequency above or below the integration range of f=[0.12,0.25] of our 1/f test, and so may lead to an incorrect failure to reject the null. We examined the power of the 1/f test for the subset of simulations where the resulting subclone does lie within the integration range (Figure 1I). We found that in the large proportion of cases, the 1/f test correctly rejected neutrality, although the challenge of hypermutant subclonal tails partially remained. Therefore, if the signature of selection lies within the integration range, the 1/f test correctly classifies neutral vs non-neutral tumours. Consequently, contrary to Tarabachi and colleagues’ assertions, effectively neutral dynamics imply a 1/f distribution, and *vice versa*. We again do acknowledge that the choice of integration range is a weakness of the 1/f analytical test, and that if a subclone lies away from the integration range (either it is very small or almost fixed within the population), neutrality may not be rejected. That is the reason we have now developed better methods that use the entire VAF distribution^3^.

Importantly, we are pleased that in their letter, Tarabichi and colleagues confirm that both in the presence of a purely neutral process and in the case of subclonal selection, 1/f neutral tails are predicted to be pervasive in cancer data. In some of their simulations, they are so pervasive that they dominate the entire VAF distribution. This is because 1/f tails are a simple consequence of clonal growth, with each individual clone generating its own neutral tail during the expansion^23^. Critically, the large majority of current subclonal reconstruction studies do not account for 1/f tails, and may have erroneously classified 1/f tails as subclones, a problem that we recently highlighted^4^. The results from Tarabichi *et al*. therefore corroborate our previous findings, and prompt the need for the reanalysis of multiple cancer genomic studies to account for this major confounding factor.

## 4. Analysis of subclonal selection using dN/dS ratios

Using a test inspired by the classical dN/dS method, the authors claim to find evidence of subclonal selection in cancers classified as neutral with our 1/f test. Specifically, they first classify the tumours as neutral/non-neutral using our 1/f method, and then pool the subclonal mutations in 192 known cancer genes from different tumours to calculate a dN/dS value for neutral vs non-neutral groups. Subclonal mutations in the neutral group should not contain evidence of selection (dN/dS should not be significantly higher than 1). Conversely, subclonal mutations in non-neutral cancers, as well as clonal mutations in all cancers are expected to contain selected genes (even in neutrally growing tumours, selection was present during tumorigenesis), thus leading to dN/dS>1. Tarabichi and colleagues claim they find significantly positive dN/dS values for subclonal mutations in neutral tumours, thus contradicting our results.

In the attempt to address the Tarabichi *et al*. criticism, we have reproduced their analysis using their dN/dS calculation method^13^. Since the authors did not mention which 192 cancer genes they used, we considered the set of 369 cancer genes from Martincorena *et al*. 2017^13^. We applied their method to the cohort of colorectal and gastric cancers analysed in our original manuscript (Figure 2A and B). For the pan-cancer analysis, we could not fully reproduce their plots because the CAVEMAN somatic calls the authors used in the letter and in their dN/dS manuscript^13^ are not publically available. We therefore reanalysed the pan-cancer TCGA dataset using the variant calls publically available through the GDC data portal (see Methods for details and Figure 2C).

**Figure 2.**
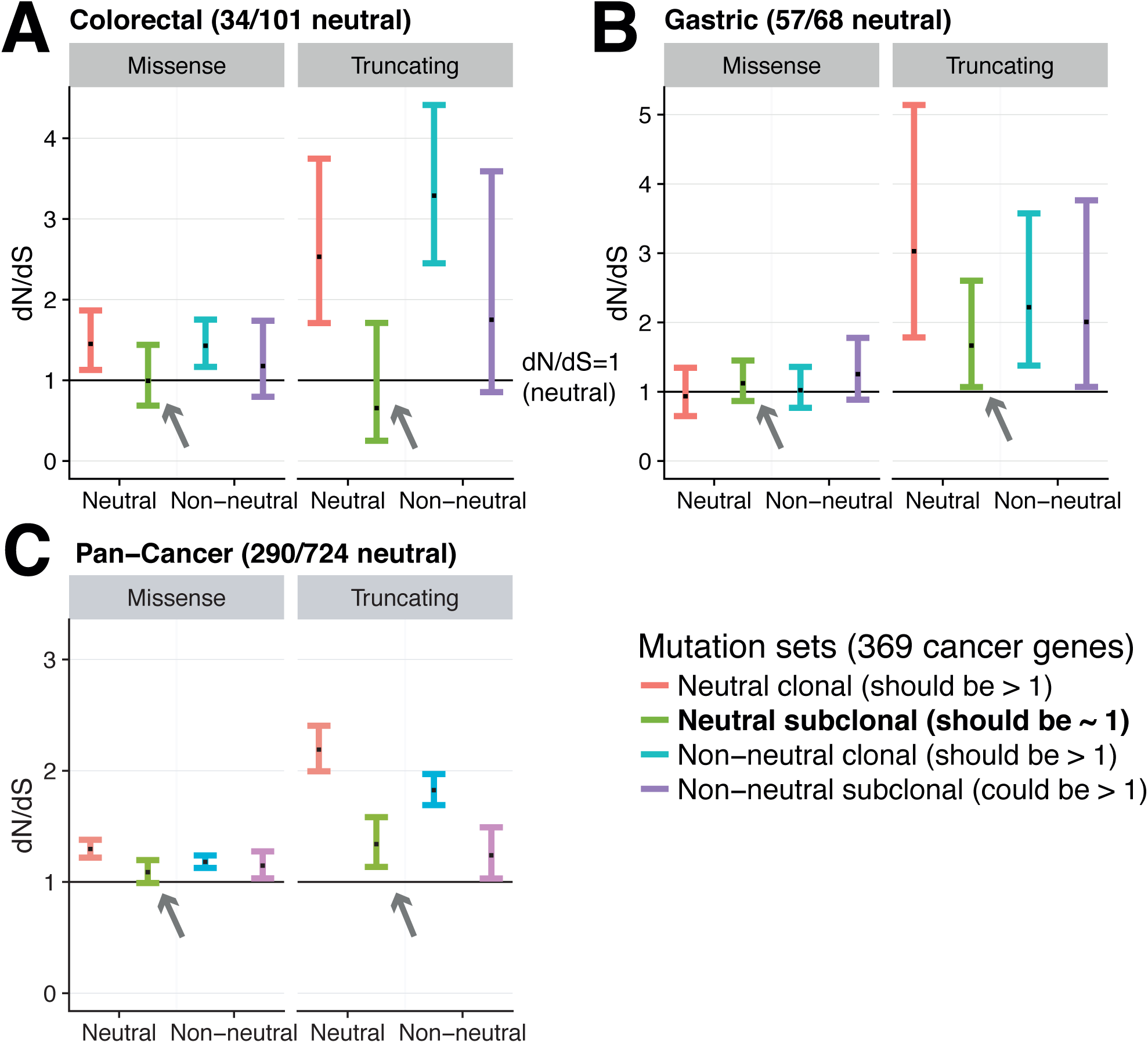
Detecting subclonal selection with dN/dS analysis. dN/dS analysis using Martincorena *et al*. 2017 method applied to the colorectal cancers (panel **A**) and gastric cancers we analysed in our original Williams *et al*. 2016 manuscript (from ref^20^ panel **B**), as well as TCGA pan-cancer analysis using newly available GDC calls to reproduce Tarabichi’s dN/dS pan-cancer analysis (panel **C**). In each type of cancers, the cancers were classified as neutral or non-neutral using the 1/f test, and the dN/dS values of clonal and subclonal variants assessed using the Martincorena method for the pooled variants in each group.

We found that subclonal missense mutations in neutral-classified cancers, precisely as predicted by neutrality, had dN/dS values that were not significantly different from 1 in all cohorts, thus corroborating our findings (Figure 2, arrows). This recapitulates Tarabichi’s analysis, which also shows that subclonal missense mutations in neutral-classified cancers have a dN/dS not significantly different from 1 (i.e. does not reject neutrality). We also note that both our and Tarabichi’s analysis does not consistently yield dN/dS>1 in cases where tumours are classified as non-neutral. Therefore, for missense mutations, dN/dS estimates are consistent with our orthogonal method for detecting selection based on VAF distributions.

We do note that subclonal truncating mutations in neutral-classified gastric and pan-cancer cohorts were above 1, although the confidence interval was not dramatically higher. It is important to appreciate that dN/dS values from truncating variants are extremely noisy because of the tiny number of these mutations in cancer cohorts (i.e. note large confidence intervals for truncating with respect to missense). We then manually inspected the subclonal truncating mutations in neutral cancers from the pan-cancer set (Figure 2C) by comparing them with the genes reported as drivers in each tumour type in the INTOGEN database (https://www.intogen.org). We found little overlap, with only 2 TP53 mutations in LGG and LIHC and 2 ARHGAP35 and 3 DICER1 mutations in UCEC and 1 FAT1 mutation in HNSC that were suggested to be pathogenic (8 cases out of 290 neutral tumours). Furthermore, some of these mutations were found in high copy number regions where we would expect the other copies to compensate for any loss of function (see Figure S1). We do acknowledge that a small number of these cases could have been misclassified as neutral with this original version of the 1/f test.

In general, we notice that for truncating mutations, the set of variants for which it is hardest to estimate the expected background mutation rate (because there are few – and hence the extremely large error bars), the tool the authors employ seem to consistently report large values of dN/dS. This phenomenon seems to occur in our dataset, as well as in the Martincorena *et al*. 2017^13^ study (Figure 1B in the paper), where instead missense mutations produce reasonable values around 1 or slightly higher. Even in normal tissue, where truncating mutations (presumably in tumour suppressor genes) are not in regions of loss of heterozygosity and therefore theoretically non-functional, the values of dN/dS are reported to be even higher than in cancer (Figure 1C in Martincorena *et al*. 2017). We cannot exclude therefore, that the values of dN/dS for truncating mutations may suffer a systematic positive bias. Together this makes the interpretation of the dN/dS of truncating mutations more difficult than missense mutations.

Moreover, caution is warranted in interpreting dN/dS ratios from *subclonal* mutations. Specifically, dN/dS estimates were originally developed in comparative genomics to quantify selection pressures on genetic loci using fixed substitutions in divergent species and not mutations in closely related lineages such as subclonal variants in cancer. In this scenario, the population genetic processes of fixation and loss become important in the interpretation of dN/dS. One consequence of this is that the simple monotonic relationship between the selection coefficient and dN/dS values is lost^14^. Another complicating factor is the interpretation of dN/dS in growing populations, where selection is inefficient and consequently insufficient time may have elapsed to enrich or deplete selected mutations, which is particularly relevant for subclonal dN/dS values measured in an expanding neoplasm. Whereas dN/dS ratios are a very powerful and important tool to analyse clonal mutations in cancer, as some of the authors recently demonstrated^13^, their interpretation for subclonal mutations is still debatable. Indeed, the dN/dS approach the authors used^13,15^ has already been disputed in the context of subclonal mutations^16^, with neutrally evolving subclones producing positive dN/dS. This is a paradox that remains unsolved^17^.

Despite some disagreements, Tarabachi and colleagues provided some valid constructive criticism of our original manuscript. In our assessment however, none of these points of criticism invalidate our original conclusion that neutral evolution provides an entirely adequate description of the pattern of intra-tumour heterogeneity that has been observed to date across many tumours.

## Methods

For the simulations in Figure 1, we used a classical stochastic branching process approach, equivalent to the one we recently presented^3^ and virtually identical to the one used in the Tarabichi *et al*. letter. Specifically, we use: death rate of 0.2 (parameter ‘p(cell_death)’), birth rate of 1 (parameter *λ*) fitness advantage of new subclone adv_subclone_=[0,1], Poisson distributed background mutation rate with mean *μ* = 16, mutation rate of new subclone *μ_subclone_*=[2, 1024], introduction of new subclone at the 8^th^ generation, tumour grows until 20^th^ generation, binomial sampling of the variant alleles, Poisson distributed coverage (100X depth), and a total of 200 simulations for parameter combinations. To prevent dying out of a cell type the last member was never removed. We have provided our simulation code, as well as a script to regenerate the simulations in Figure 1A as Supplementary Information.

To reproduce the Tarabichi *et al*. pan-cancer dN/dS analysis, we downloaded 8,455 TCGA tumours from different cancer types from the GDC portal (https://portal.gdc.cancer.gov/) and selected cancers with high purity (>=70%) using variants called with Mutect2, as we did in our original manuscript. We then adjusted the VAF for the purity obtained from ref^18^ and classified tumours as neutral/non-neutral using our 1/f test on diploid regions^19^, employing the f=[0.12,0.24] integration range and requiring a minimum of 12 subclonal mutations, again as in our original manuscript. Out of the 8,455 tumours, 724 satisfied all the conditions (see Table S2). We then run the method dndscv described in Martincorena *et al*. 2017^13^ in the four datasets (neutral clonal, non-neutral clonal, neutral subclonal and non-neutral subclonal). We classified a mutation subclonal if its adjusted VAF was <=0.24 to be consistent with the 1/f integration range. We must note that the newly published ploidy estimates^19^ improved the classification, with 40% of neutral cases identified in this cohort (290/724) with respect to the 31% in our original analysis. We frankly acknowledge the limitations of the 1/f test integration range, which remains suboptimal but we kept the same as before for consistency with the original manuscript and to reproduce the Tarabichi *et al*. analysis.

**Figure S1.** VAF distribution of TCGA pan-cancer neutral cancers with subclonal truncating mutations. For each neutral cancer from the TCGA pan-cancer cohort in which a truncating subclonal mutation in one of the 12 cancer genes with a significant elevated dN/dS value was present, here we plot the VAF distribution adjusted for purity and highlight the frequency of the truncating variant. Whereas the majority of variants lied on the 1/f tail and were suggested not to be pathogenic following INTOGEN database analysis, we do acknowledge that a handful may have been cancers miscalled as neutral.

**Table S1.** List of TCGA pan-cancer cases selected for neutral evolution analysis. Out of 8,455 initial cases available through GDC, a final set of 724 was high purity (>=70%) and had at least 12 subclonal variants in diploid regions. The proportion of neutral cases identified was 40%.

